# Ergosterol-depleted clinical isolates of *Nakaseomyces glabratus* can develop multi-drug resistance without apparent fitness and virulence defects

**DOI:** 10.1101/2025.08.28.672802

**Authors:** Alexander M. Aldejohann, Nadja Thielemann, Aina Martinez Zurita, Christoph Müller, Tom Gräfenhan, Richard Kriz, Heimo Lagler, Isabell S. Behr, Nathalie Reus, Annika Schöninger, Grit Walther, Lena-Marie Mazan, Hannah Wilhelm, Birgit Willinger, Christina A. Cuomo, Oliver Kurzai, Ronny Martin

## Abstract

**Objectives:** *Nakaseomyces glabratus* (formerly *Candida glabrata*) is a leading cause of invasive candidiasis and rapidly develops antifungal drug resistance during treatment. An increasing number of clinical isolates shows reduced susceptibility to echinocandins and azoles, leaving amphotericin B (AMB) as a last therapeutic option. Resistance of *N. glabratus* to this drug is rare and its underlying mechanisms are still not fully understood. Here, we describe two independent multidrug resistant (MDR) bloodstream isolates displaying resistance to AMB and anidulafungin (ANF) as well as a reduced susceptibility to azoles.

**Methods:** Whole-genome sequencing and sterol profiling were performed on nine clinical *N. glabratus* isolates which were resistant to ANF and displayed resistance or low susceptibility to fluconazole (FLU) and AMB. The transcriptional response of reference strain CBS138 and an AMB^R^+ANF^R^ isolate was analyzed by RNA-seq. Furthermore, *PDR1* was deleted in the latter isolate to examine its influence on efflux pump gene expression. Additionally, fitness and virulence of the AMB^R^+ANF^R^ isolate were examined in growth assays and a *Galleria mellonella* infection model.

**Results:** Loss of function mutations in the genes *ERG3* and *ERG4* is linked to ergosterol depletion and AMB resistance. Ergosterol depletion also contributed to a Pdr1-mediated up-regulation of *ERG* and ABC transporter genes which was associated with low FLU susceptibility. The AMB^R^ isolates displayed no fitness defects and one of them was fully virulent in a *G. mellonella* infection model.

**Conclusions:** These findings demonstrate that ergosterol depletion in *N. glabratus* leads to AMB resistance without affecting fitness or virulence.

## Introduction

Invasive candidiasis is a life-threatening fungal infection, caused by yeasts including *Candida albicans* and *Nakaseomyces glabratus*, *and* comprises bloodstream infections, but also dissemination to organs like liver and kidney, accounting for approximately 1.5 million cases per year with attributable mortality of up to 60% [1,2]. While *C. albicans* is the most virulent and best studied pathogen and still accounts for approximately 50% of systemic *Candida* infections, non-*albicans Candida* species are of increasing importance as they often develop antifungal drug resistance or even MDR [1,4]. The recommended first-line treatment for systemic *Candida* infections are echinocandins [5,6]. They bind the catalytic subunit of the ß-1,3-D-glucan synthetase and inhibit the ß-1,3-D-glucan biosynthesis, leading to disruption of cell wall integrity and osmotic imbalance [7]. Echinocandin resistance is mainly caused by point mutations in the hot spot regions of the *FKS* genes which encode the enzyme’s catalytic subunit [8]. Echinocandin resistance in *N. glabratus* is rare but increased in recent years [9]. Due to the intrinsically low susceptibility of *N. glabratus* to azoles and a high proportion of azole-resistant isolates, acquisition of echinocandin resistance often results in MDR, leaving liposomal AMB as an indispensable option, despite severe side effects for the patients such as high nephrotoxicity or serum electrolyte changes [1,5,6]. AMB binding to ergosterol leads to either formation of small ion channels or ergosterol extraction from the cell membrane [10–12]. AMB resistance is still very rare among *Candida* species. Ergosterol depletion seems to be the major resistance mechanism but is often linked to high fitness costs [7,13,14].

Here, we describe AMB resistance in two independent clinical *N. glabratus* isolates and show that combined mutations in *ERG3* and *ERG4* are responsible for the resistance but do not result in fitness defects. One of the isolates also displayed decreased susceptibility to anidulafungin and azoles. The latter was Pdr1-dependent and likely triggered by ergosterol depletion. Thus, ergosterol depletion in the context of AMB resistance can directly result in MDR phenotypes in *N. glabratus*.

## Material and Methods

### Strains and media

All strains used in this study are listed in Table S1. They were routinely grown in YPD medium (20g/L glucose, 10g/L yeast extract, 20g/L peptone) at 37°C unless otherwise indicated.

### Antifungal drug susceptibility testing (AFST)

AFST was either performed with EUCAST-based broth microdilution [15] or with Etests (Biomérieux) according to manufacturer’s instructions.

### Plasmid and strain construction

The procedures of plasmid and strain construction can be found in the supplement (Data S4). Oligonucleotide primers and plasmids are listed in Table S2 and S3.

### DNA Isolation and whole-genome sequencing (WGS)

The ZR Fungal/Bacterial DNA MiniPrep kit (Zymo Research, Irvine, CA, USA) was used to extract fungal genomic DNA. Genomic libraries were constructed and barcoded using the NEBNext Ultra DNA Library Prep kit for Illumina (New England Biolabs, Ipswich, MA, USA) and then sequenced using the Illumina platform. Further details of the data analysis are available in the supplement (Data S4).

### Measurement of sterol components

The sterol composition of clinical *N. glabratus* isolates was determined by gas chromatography-mass spectrometry (GC-MS) as previously described [16].

### Cell wall composition analysis

Details of the staining and measurement of *N. glabratus* cell wall components by flow cytometry are listed in the supplement (Data S4).

### Transcriptome analysis

Fungal cells were grown overnight in YPD at 37°C. 1×10^6^ cells / mL were then added to prewarmed YPD with or without 1 µg/mL AMB and grown for 1 h at 37°C. Cells were harvested by centrifugation and total RNA was isolated as previously described [17]. Library preparation and RNA-sequencing was performed by the Core Unit SysMed Würzburg. Details of the data analysis are available in the supplement (Data S4).

### Data availability

Sequencing data are available in the National Center for Biotechnology (NCBI) Sequence Read Archive (SRA) under BioProject PRJNA1299776.

### Gene expression analysis

100 ng/µl total RNA from the same conditions as used for the transcriptome analysis were the template for RT-qPCR using the Luna Universal One-Step RT-qPCR Kit with SYBR Green (New England Biolabs). Table S2 lists all used oligonucleotide primers. Gene expression was calculated with the ΔΔCt method [18]. *RDN5.8* and a control RNA (5h YPD, 37°C) were used for normalization. Data from independent biological triplicates were compared with a two-tailed, unpaired student’s *t*-test and p values ≤ 0.05 were regarded as statistically significant.

### Assessment of *in vivo* pathogenicity in *Galleria mellonella* larvae

Prior to the inoculation with *N. glabratus*, *Galleria mellonella* larvae weighing 220-280 mg fasted for 24 hours. Larvae were wiped with 70% ethanol, clustered into groups of 15 and placed in individual petri dishes for observation. *N. glabratus* CBS138 and NRZ-2016-252 were grown overnight at 37°C on Sabouraud dextrose agar. Fungal cells were then harvested and suspended in sterile phosphate-buffered saline (PBS). Each larva within a group was injected with 10 µL of the yeast cell suspension with 2×10^9^ colony-forming units /mL, resulting in the inoculation of 2×10^7^ yeast cells into each larva. The control group larvae were injected with sterile PBS. The infected larvae were incubated at 37°C. Larvae of the intervention groups were injected with AMB (5 mg / kg body weight) one hour after inoculation. Larval survival, indicated by melanisation and mobility, was periodically monitored during the next seven days. Survival rates were visualized using Kaplan-Meier plots.

## Results

### *ERG3* mutations are enriched among MDR isolates

We have previously analyzed the emergence of echinocandin-resistant clinical isolates of *N. glabratus* in Germany [9]. Based on this work, we further examined strains, which displayed (i) resistance to the echinocandin ANF without harboring *FKS* hot spot mutations and (ii) additional resistance to either FLU or AMB (=MDR isolates, Figure 1 A). Whole-genome sequencing was performed for all six identified MDR isolates and three isolates with isolated ANF resistance (ANF^R^) and one control isolate (AMB^S^, ANF^S^, FLU^I^). The MDR strains formed no clear cluster but showed an enrichment of *ERG3* mutations (Figure 1 A, B). The two AMB^R^ strains were genetically unrelated. Their respective closest relatives were AMB^S^ and displayed a high genetic variation compared to the AMB^R^ strains (Figure 1 B, C). We identified putative loss of function mutations in the *ERG3* genes of the strains NRZ-2017-099 (M1*), NRZ-2016-252 (Q26*) and NRZ-2016-150 (K133^del^) (Figure 1 A). The latter two strains also displayed putative loss of function mutations in the *ERG4* genes: T158^fs^ and Y327* (Figure 1 A). Despite identifying several *FKS* mutations in our strains, only the isolate NRZ-2017-099 harbored two mutations which might explain ANF resistance: K1323N in *FKS1* and T970fs in *FKS2* (Figure 1 A). Despite displaying ANF resistance, no *FKS* mutations were identified in NRZ-2016-252, NRZ-2016-191 and NRZ-2017-475 (Figure 1 A).

**Figure 1.**
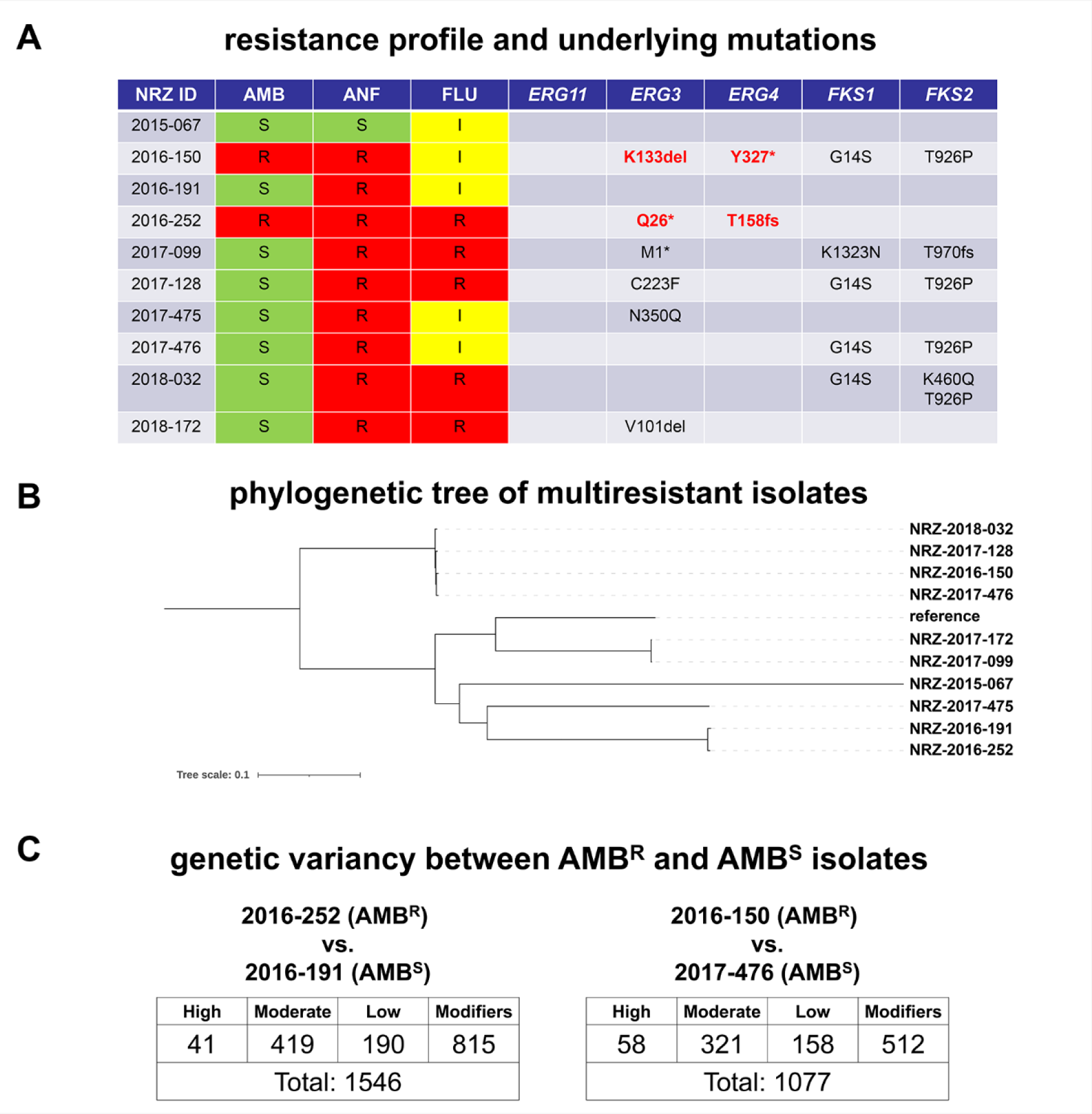
Resistance profile and genetic characteristics of MDR *N. glabratus* isolates. (A). Summary of antifungal susceptibility testing (AFST) and underlying mutations in possible resistance genes. AFST was performed with broth microdilution and based on EUCAST clinical breakpoints, the isolates were determined as susceptible (S), increased-dose-dependent (I) or resistant (R). Shown mutations in the indicated genes were identified by whole genome sequencing (WGS). (B) WGS-based phylogenetic tree of the examined *N. glabratus isolates*. (C) Genetic diversity of AMB^R^ isolates NRZ-2016-252 and NRZ-2016-150 and their closest relatives NRZ-2016-191 and NRZ-2017-476.

### Cell wall composition of MDR isolates

Therefore, we used flow cytometry to measure the amounts of chitin, glucan and mannan within the isolates as an altered composition is sometimes associated with echinocandin resistance. There was no clear link to ANF resistance, but the AMB^R^+ANF^R^ isolate NRZ-2016-252 and the ANF^R^ strain NRZ-2016-191 which both harbor no *FKS* mutation, tend to have more of chitin, glucose and mannan (Figure S1).

### Ergosterol depletion correlates with resistance to AMB

We additionally examined the sterol composition of the strains which exhibited resistance to AMB or azoles. With a percentage of 77%, ergosterol was the main sterol in the reference strain CBS138 (Figure 2 A, Table S5). The *ERG3* mutations M1*, V101del and C223F in the isolates NRZ-2017-099, NRZ-2017-128 and NRZ-2018-172 were associated with decreased ergosterol levels and the accumulation of ergosta-7,22-dien-3β-ol and ergost-7-en-3β-ol, indicating that the function of the sterol C5-desaturase Erg3 is disturbed or lost (Figure 2 A). The AMB^R^ + ANF^R^ isolates NRZ-2016-252 and NRZ-2016-150 had extremely low ergosterol concentrations (0.1-1%) while ergosta-7,22,24(28)-trien-3β-ol increased up to 85% (Figure 2, Table S5). This is only possible if neither Erg3 nor Erg4 work properly, fitting to the identified mutations in both strains (Figures 1A and 2 B). Strains with low ergosterol concentration of 4-6% are still susceptible to AMB, indicating that a complete depletion is required for AMB resistance (Figure 2 B).

**Figure 2.**
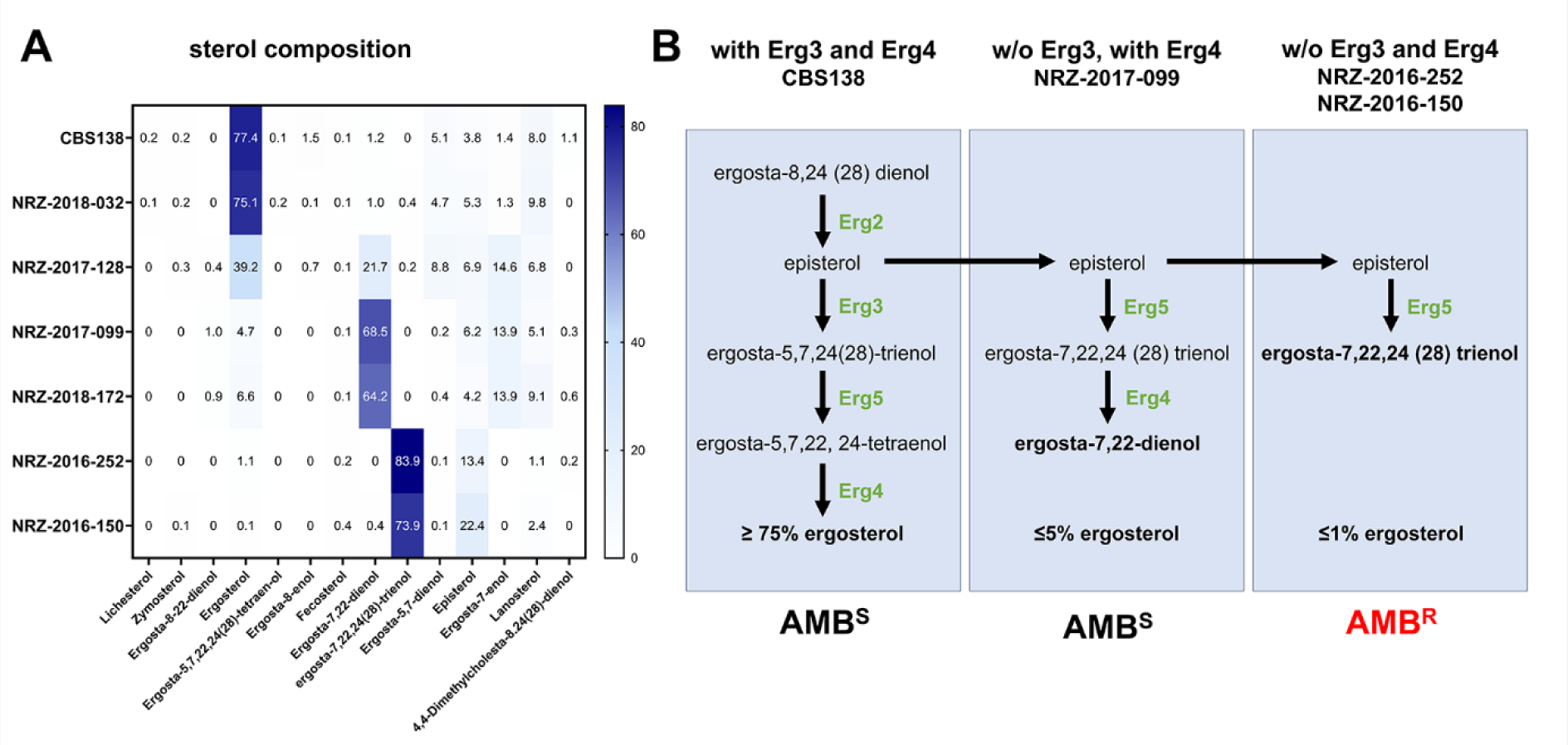
Sterol composition analysis of CBS138 and the clinical isolates. (A) Sterol composition of the indicated strains was analyzed by gas chromatography-mass spectrometry (GC-MS). Shown are the percentages of the single sterols in the indicated strains. (B) Illustration of the effects of the absence of either Erg3 and/ or Erg4 in AMB^S^ and AMB^R^ strains. Shown are branches of the ergosterol biosynthesis pathway in absence of Erg3 and/ or Erg4 leading to the production of non-physiological predecessors.

### Absence of functional *ERG3* and *ERG4* induces resistance to AMB and ANF

We hypothesized that the simultaneous loss of function in Erg3 and Erg4 enzymes cause AMB resistance and also lead to resistance against ANF and low susceptibility to FLU. To confirm this, we replaced the *ERG3*^Q26*^ allele in NRZ-2016-252 with the wild type allele of CBS138. The resulting mutant became susceptible to AMB and ANF (Figure 3 A). Surprisingly, the strain was FLU^R^, maybe caused by the still present *ERG4*^T158fs^ mutation (Figure 3 A). Additionally, we deleted *ERG3* and *ERG4* in the background strain HTL. The resulting double mutant displayed AMB and ANF resistance and a low susceptibility to FLU, similar to NRZ-2016-252 (Figure 3 B).

**Figure 3.**
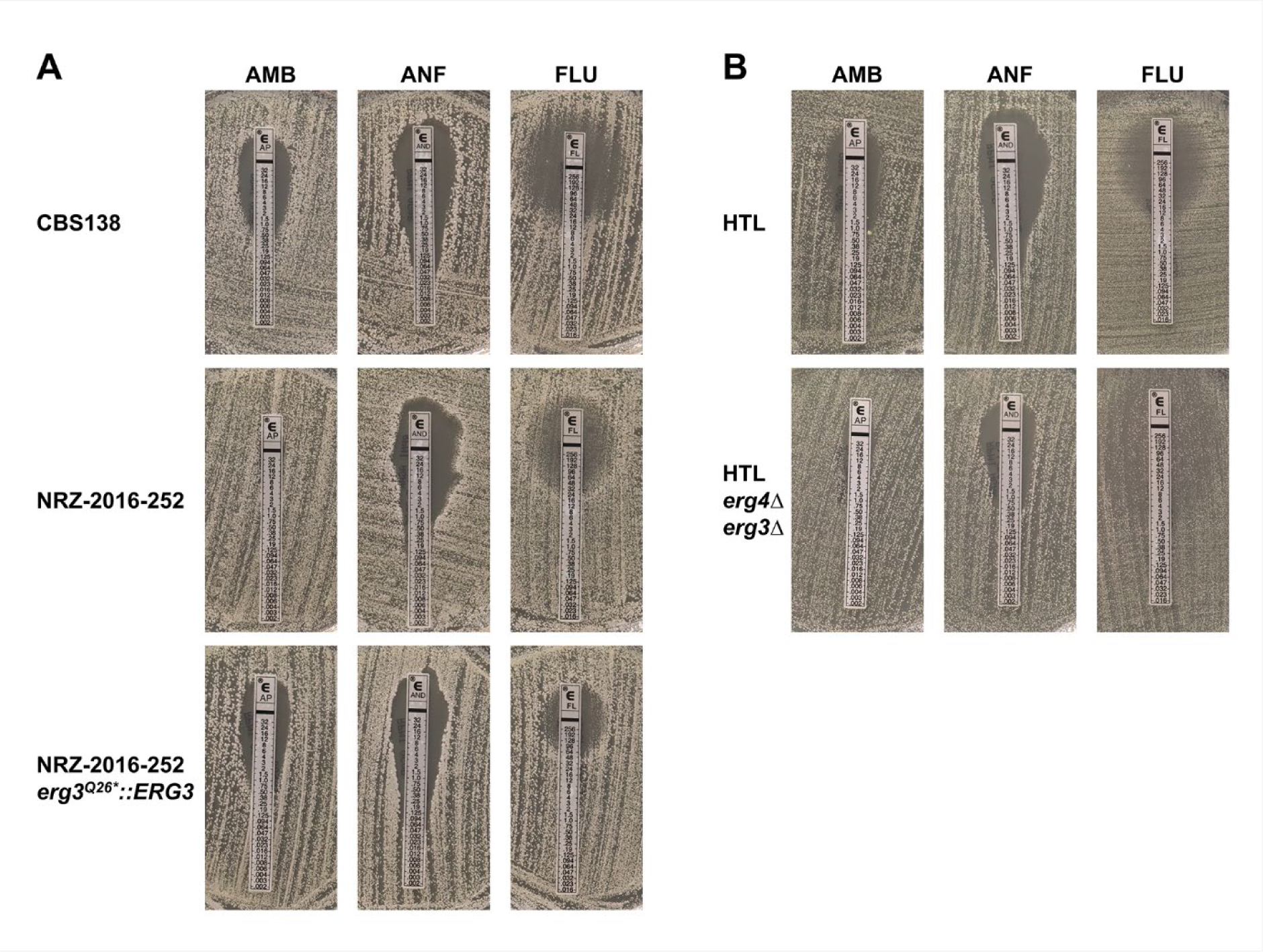
Absence of *ERG3* and *ERG4* is required for resistance to AMB and ANF. (A) Integration of a wild type *ERG3* allele into the AMB^R^ isolate NRZ-2016-252 increased susceptibility to AMB and ANF but not to FLU. The indicated strains were plated onto RPMI1640 medium and E-test stripes for AMB, ANF and FLU were applied. The plates were then grown for 48h at 37°C before pictures were taken. (B) *ERG4* and *ERG3* were deleted in the *N. glabratus* HTL background strain. HTL and the *erg4*Δ/ *erg3*Δ double mutants were plated and antifungal drug susceptibility was determined with E-tests as described in (A).

### Pdr1 links ergosterol depletion and low susceptibility to FLU

To better understand the fungal cell response to AMB, we examined the transcriptomes of the AMB^S^ strain CBS138 and the AMB^R^ isolate NRZ-2016-252 after 1 h incubation in YPD with or without 1 µg/ml AMB at 37°C. 1203 genes were differentially expressed in CBS138 in response to AMB (Figure 4 A, dataset S6). Interestingly, 68% of them were differentially expressed in the AMB^R^ strain NRZ-2016-252 in absence of AMB, indicating that this strain is already well-adapted to AMB (Figure 4 A, dataset S6). Among the upregulated genes in the AMB^R^ strain were stress-related genes (*UPC2B, TYE7*, *ICL1*, *ICL2*, *HSP12, RAD27*), efflux pumps (*CDR1*, *PDH1*, *FLR1* and *FLR2)* and ergosterol biosynthesis genes (Figure 4 B and C, dataset S6). The two latter groups are known targets of the transcription factor Pdr1 [19,20]. Especially the up-regulation of the efflux pump genes might explain the low FLU susceptibility of NRZ-2016-252. Deletion of the *PDR1* gene caused a dramatic decline of *CDR1, FLR1* and *PDH1* transcription in NRZ-2016-252 (Figure 5 A). Compared to NRZ-2016-252, the *pdr1*Δ derivate was extremely susceptible to fluconazole, indicating that up regulation of *CDR1* and *PDH1* in NRZ-2016-252 was required for the low FLU susceptibility of NRZ-2016-252 (Figure 5 B). NRZ-2016-252 *pdr1*Δ remained AMB and ANF resistant, illustrating that Pdr1 was not required for resistance to these drugs (Figure 5 B).

**Figure 4.**
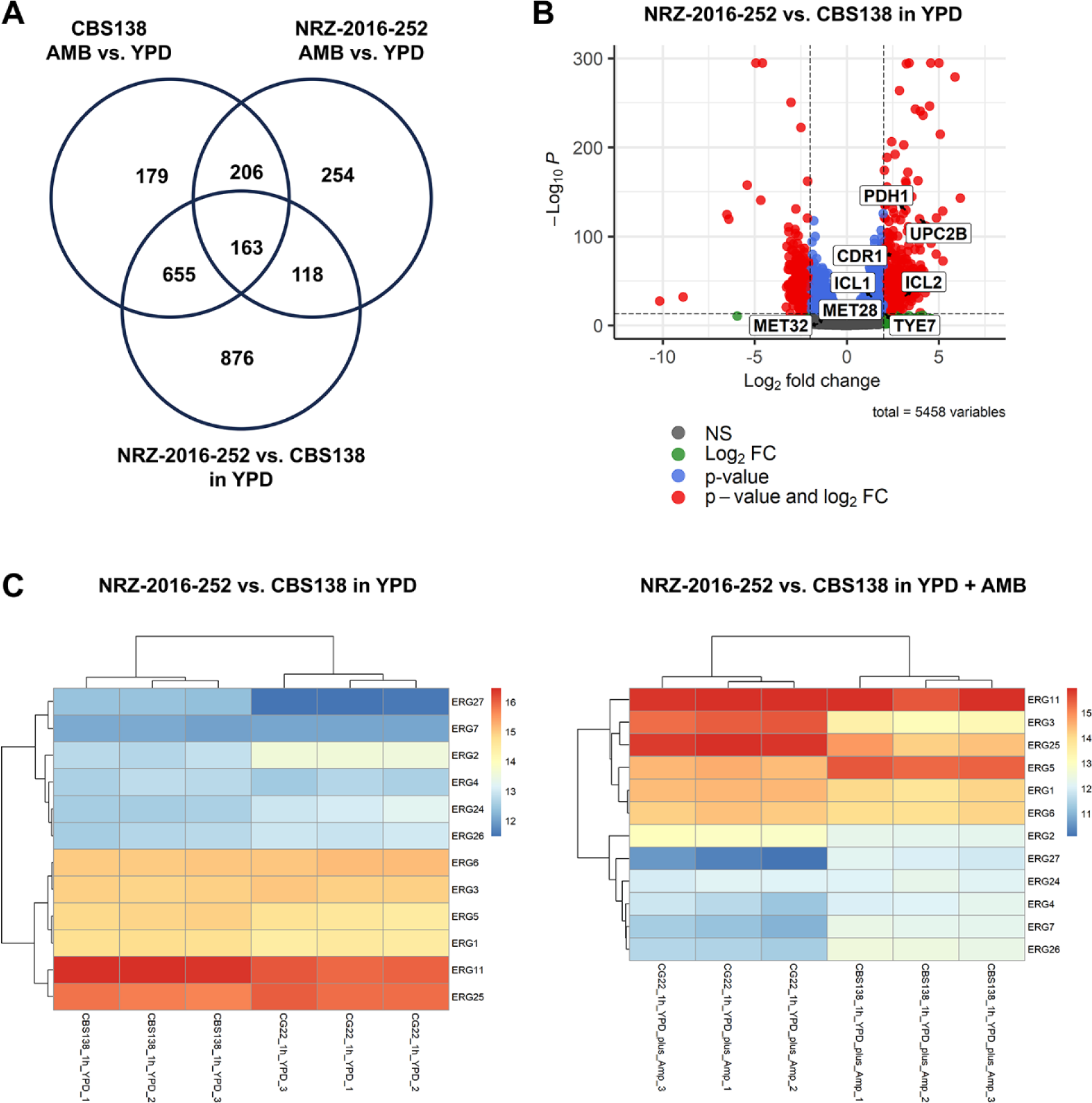
Transcriptional response of AMB^S^ and AMB^R^ strains to AMB. The AMB^S^ strain CBS138 and the AMB^R^ strain NRZ-2016-252 were grown for 1h in YPD with 1 µg/ml AMB (AMB) or without 1 µg/ml AMB (YPD) prior to RNA isolation. (A) Comparison of upregulated genes in CBS138 and NRZ-2016-252 in AMB and of genes upregulated in NRZ-2016-252 compared to CBS138 in YPD. Example genes up-regulated in NRZ-2016-252 compared to CBS138 in presence of absence of AMB are shown in the box. (B) Vulcano plot of differentially expressed genes in NRZ-2016-252 compared to CBS138 after 1 h growth in YPD at 37°C. (C) Comparison of *ERG* gene expression in NRZ-2016-252 and CBS138 in either YPD or YPD + AMB.

**Figure 5.**
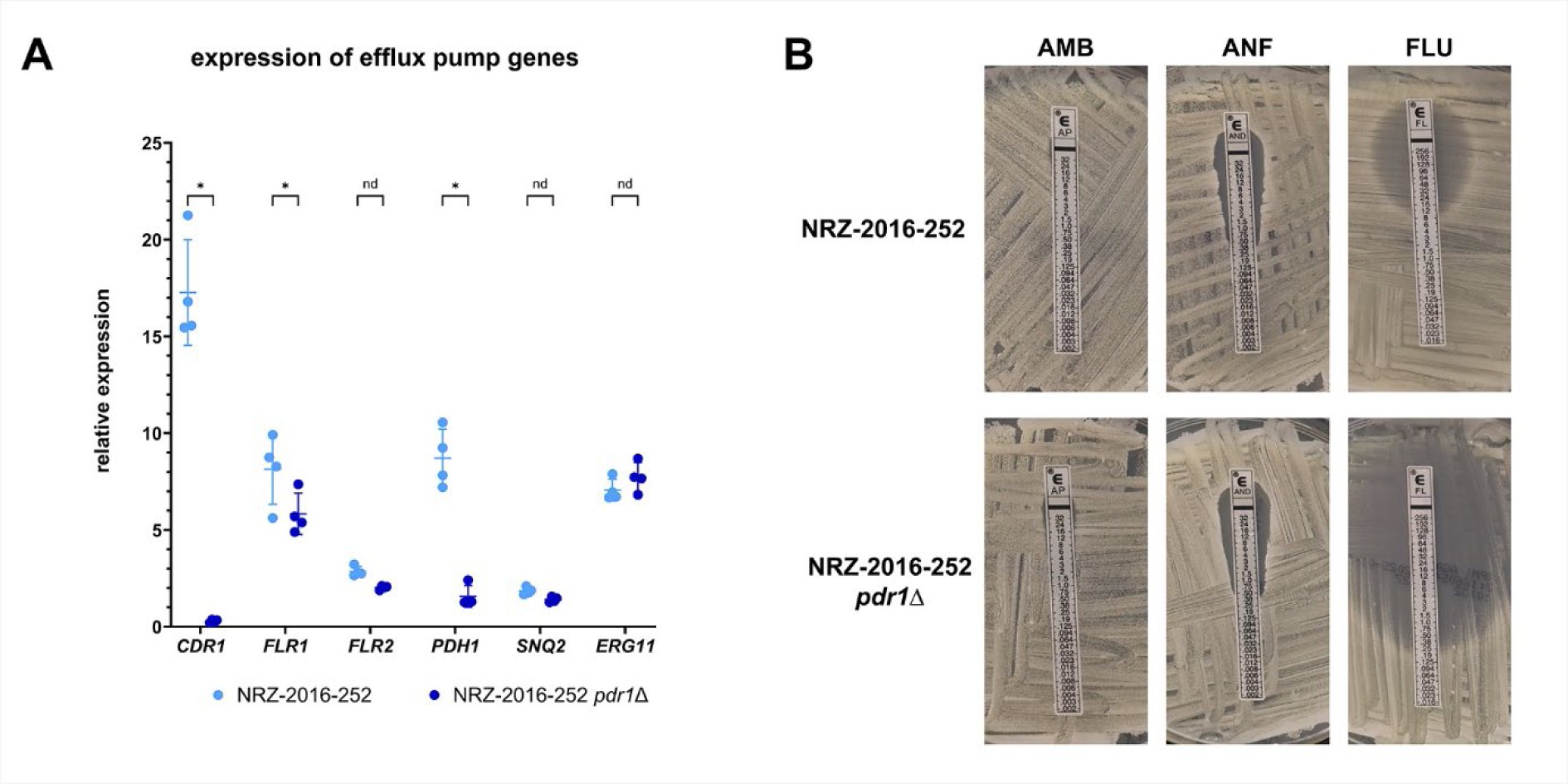
Deletion of *PDR1* led to downregulation of efflux pump genes in NRZ-2016-252 and increased susceptibility to FLU. (A) Expression of efflux pump genes in NRZ-2016-252 and NRZ-2016-252 *pdr1*Δ after 1 h growth in YPD medium at 37°C. Gene expression was normalized against CBS138 (1h YPD, 37°C) and the *RDN5.8* gene. Asterisks indicate significant changes in gene expression between the two *N. glabratus* strains (two-tailed, unpaired student’s *t*-test, p ≤ 0.05). (B) The indicated strains were plated onto RPMI1640 medium, and E-test stripes for AMB, ANF and FLU were applied. Pictures were taken after an incubation for 48h at 37°C.

### AMB^R^ isolates display no severe fitness defects or attenuated virulence

As AMB resistance is often linked to profound fitness defects, we examined the growth dynamics of the AMB^R^ isolates under different conditions. No apparent growth defects were observed in YPD medium at 37°C where both strains showed similar dynamics as the reference strain CBS138 (Figure 6 A). We then tested the growth of these strains, the isolate NRZ-2016-191 and the reference strain CBS138 under several stress conditions. Despite the ergosterol depletion, both AMB^R^ isolates showed good stress resilience at 37°C and robust growth at 42°C (Figure 6 B). Only NRZ-2016-252 displayed a growth delay under osmotic stress conditions (1.5 M NaCl) and was also more susceptible to combined stressors such as 42°C and 1.5 M NaCl or 0.0125% SDS (Figure 6 B, C).

**Figure 6.**
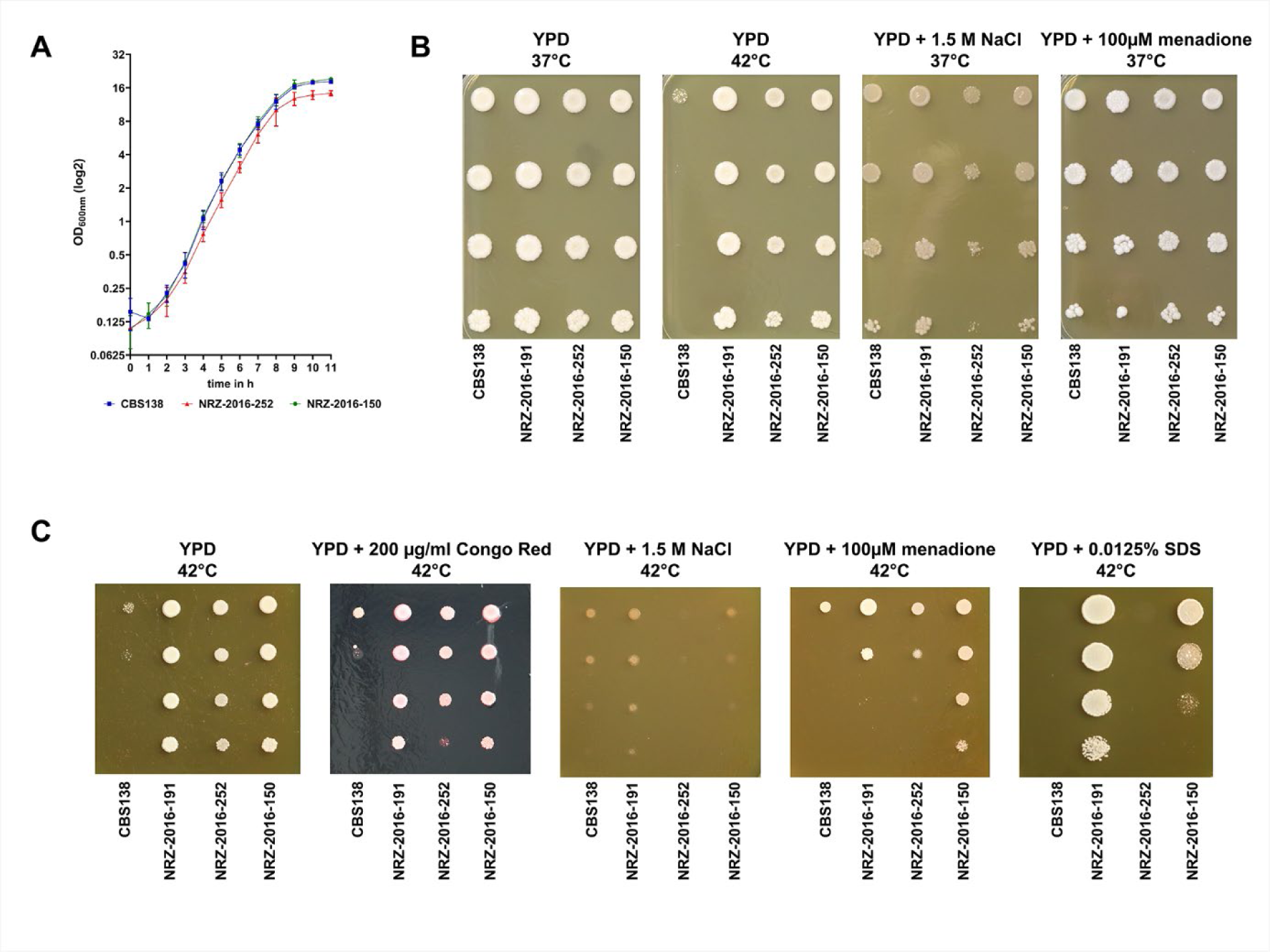
AMB^R^ isolates show no fitness defects under single but under combinatory stress conditions. (A) Growth curve of CBS138, NRZ-2016-252 and NRZ-2016-150 grown in YPD medium at 37°C for 11 h. The optical density of the three strains was measured by photometry at 600 nm. (B) The indicated *N. glabratus* strains were grown on YPD with or without 1.5 M sodium chloride or 100 µM menadione to induce osmotic and oxidate stress. The plates were incubated at 37°C for 3 days prior to photography, except one plate which was incubated at 42°C. (C) The same *N. glabratus* strains were grown on YPD at 42°C with or without 200 µg/ml Congo Red to initiate cell wall stress, 1.5 M NaCl for osmotic stress, 100 µM menadione for oxidative stress and 0.0125% SDS for cell membrane stress. The plates were grown for 3 days prior to photography.

Finally, we analyzed the virulence of the AMB^R^ strain NRZ-2016-252 in *Galleria mellonella*. The isolate displayed no attenuated virulence compared to CBS138 (Figure 7). After treatment with AMB, 86% of the larvae (12/14) survived the infection with CBS138 after 7 days (Figure 7) which was not the case for NRZ-2016-252-infected larvae which died within the first 4 days, independent from the addition of AMB (Figure 7).

**Figure 7.**
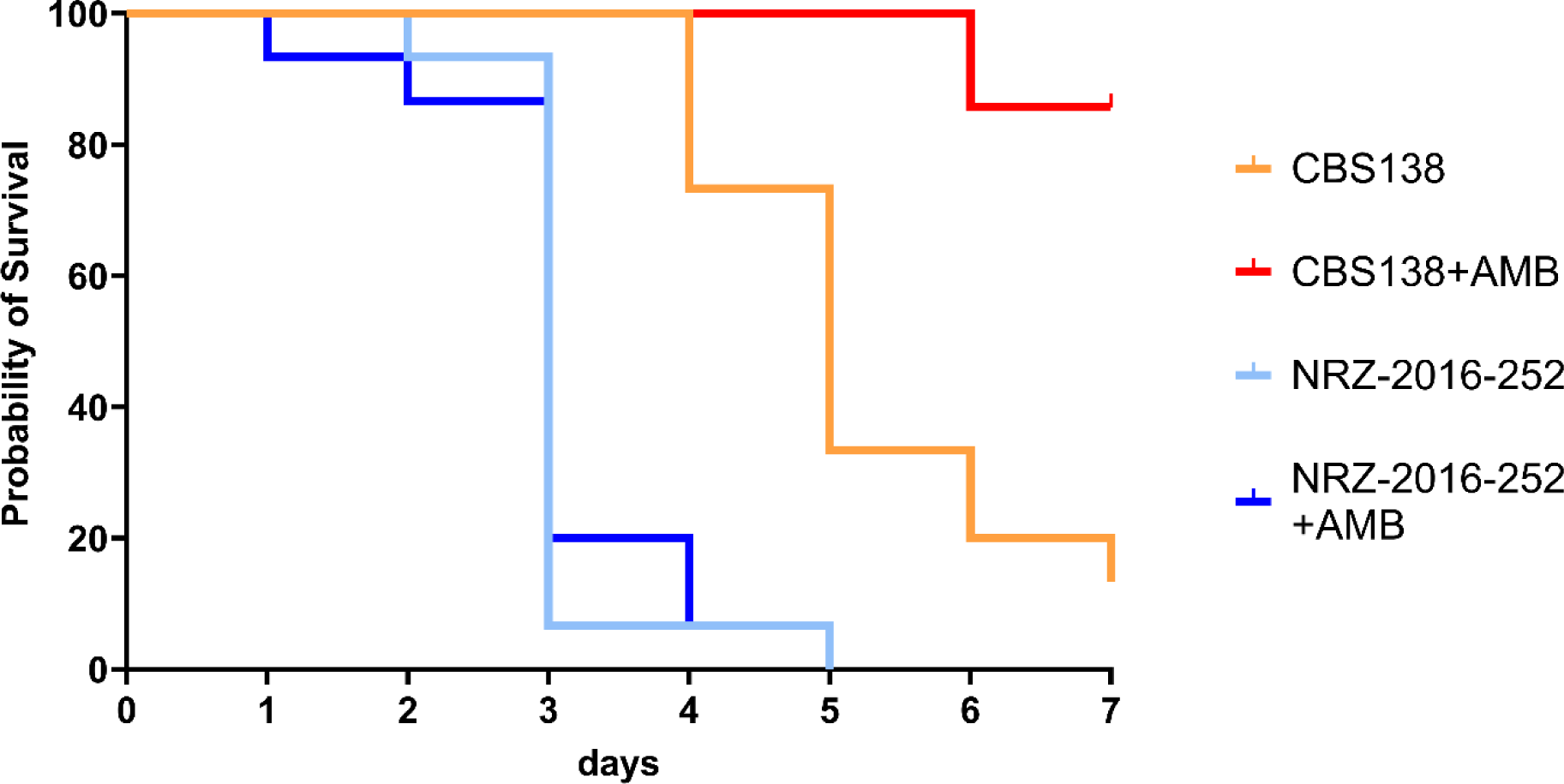
*N. glabratus* NRZ-2016-252 shows resistance against AMB under *in vivo* conditions. *Galleria mellonella* larvae were infected with 2×10^7^ *N. glabratus* cells. The infected larvae were treated with 5 mg AMB per kg body weight and incubated for up to 7 days at 37°C. The evaluation of survival of the larvae, indicated by melanisation and mobility, was first monitored 12 h post infection and then periodically each 24 h. Survival rates were visualized by Kaplan-Meier plots.

## Discussion

Our study showed that loss of function mutations in the *ERG3* and *ERG4* genes led to ergosterol depletion and consequently AMB resistance, similar to previous findings for *Clavispora lusitaniae* and *Candida auris* (syn. *Candidozyma auris*) [21,22]. Absence of functional Erg3 and Erg4 resulted in a massive shift from ergosterol to ergosta-7,22,24(28)-trien-3ßol in the two AMB^R^ isolates which was also reported for AMB^R^ *C. auris* strains [22]. This understudied sterol has a similar binding affinity to liver X receptors as ergosterol and is often enriched in ergosterol lacking fungal strains [23–25]. The primary effect of ergosterol depletion is AMB resistance as the drug can no longer bind to its target ergosterol. Additionally, it induced a transcriptional adaptation against AMB even in its absence, including the up-regulation of Pdr1-controlled genes *CDR1* and *PDH1*. The increased expression of these ABC transporter genes led to an increased efflux pump activity and therefore to reduced susceptibility to FLU which is in accordance with previous observations [19,20]. The observed association between AMB^R^ and ANF^R^ without underlying *FKS* hot spot mutations could not be fully explained. Changes in the cell wall and the cell membrane caused by the absence of ergosterol may reduce the accessibility of the ß-1,3-D glucan synthase for anidulafungin. Similar to previous works [26], our strains were susceptible to micafungin, either caused by a higher affinity or less interaction with the altered cell membrane.

Strikingly, AMB^R^ isolate NRZ-2016-252 was fully virulent in a *Galleria mellonella* infection model. Its overall *in-vitro* and *in-vivo* fitness contradicts the hypothesis that acquisition of AMB resistance is associated with high fitness costs [14, 22]. Especially, the bloodstream was previously discussed as an environment too hostile for the survival of AMB resistant strains [14], however our two AMB^R^ isolates were obtained from blood cultures. We presume that the acquisition of AMB resistance alone made the strains not more susceptible to the harsh conditions within the bloodstream. Suppressor mutations might have bypassed some defects caused by ergosterol depletion.

We described *N. glabratus* bloodstream isolates with stable AMB resistance without apparent fitness and virulence defects. In combination with the intrinsically low susceptibility to azoles and the emerging echinocandin resistance, these findings underline the threat of an increasing MDR and extensively drug resistance (XDR) in this major human fungal pathogen. A close resistance monitoring is therefore urgently needed.

## Acknowledgements

This work was supported by the German Research Foundation (DFG) through the TRR 124 FungiNet “Pathogenic fungi and their human host: Networks of Interaction”, DFG project number 210879364, project C3 (O. K.) and NIH NIAID grant U19AI110818 to the Broad Institute (C.A.C.). We want to thank Elke Huprich, Ina Gaube, Barbara Conrad, Sabrina Speiser and Margarete Göbel for their excellent technical support to this project.

## Figure Legends

**Figure S1.**
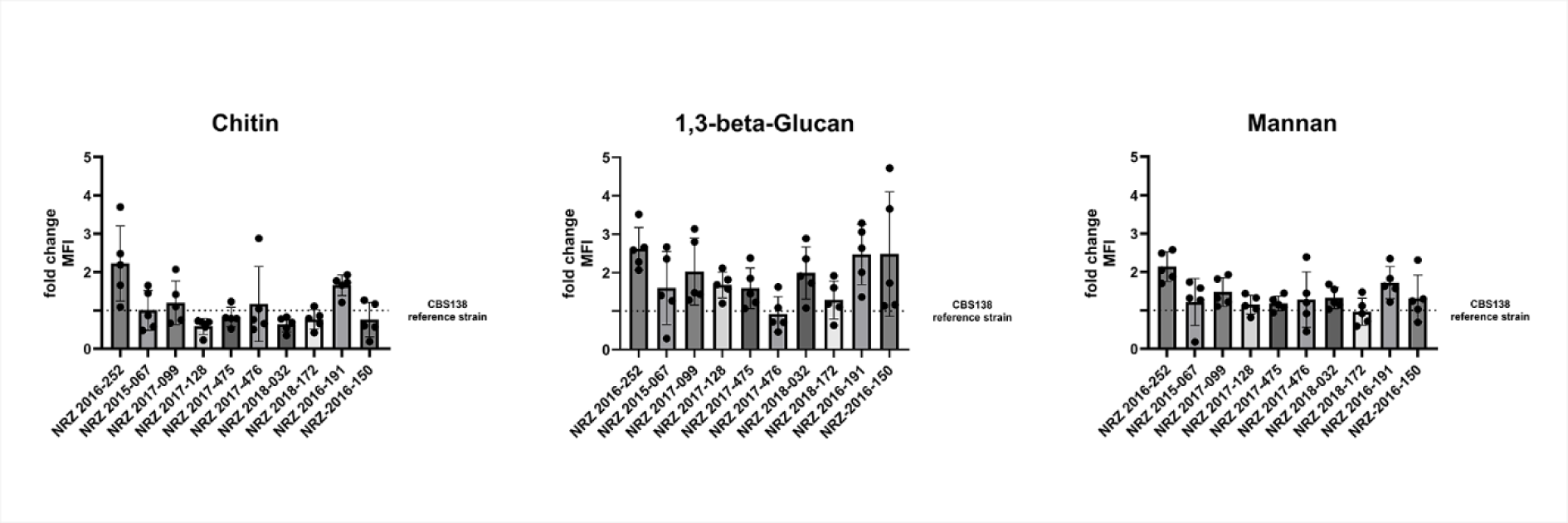
Cell wall composition of MDR *N. glabratus* isolates. The indicated *N. glabratus* strains were grown in YPD at 37°C, harvested and then stained for chitin, ß-1,3-D-glucan and mannan. The amounts of the three cell wall components were measured by flow cytometry. The values were normalized against the values of the control strain CBS138.

**Table S1.**
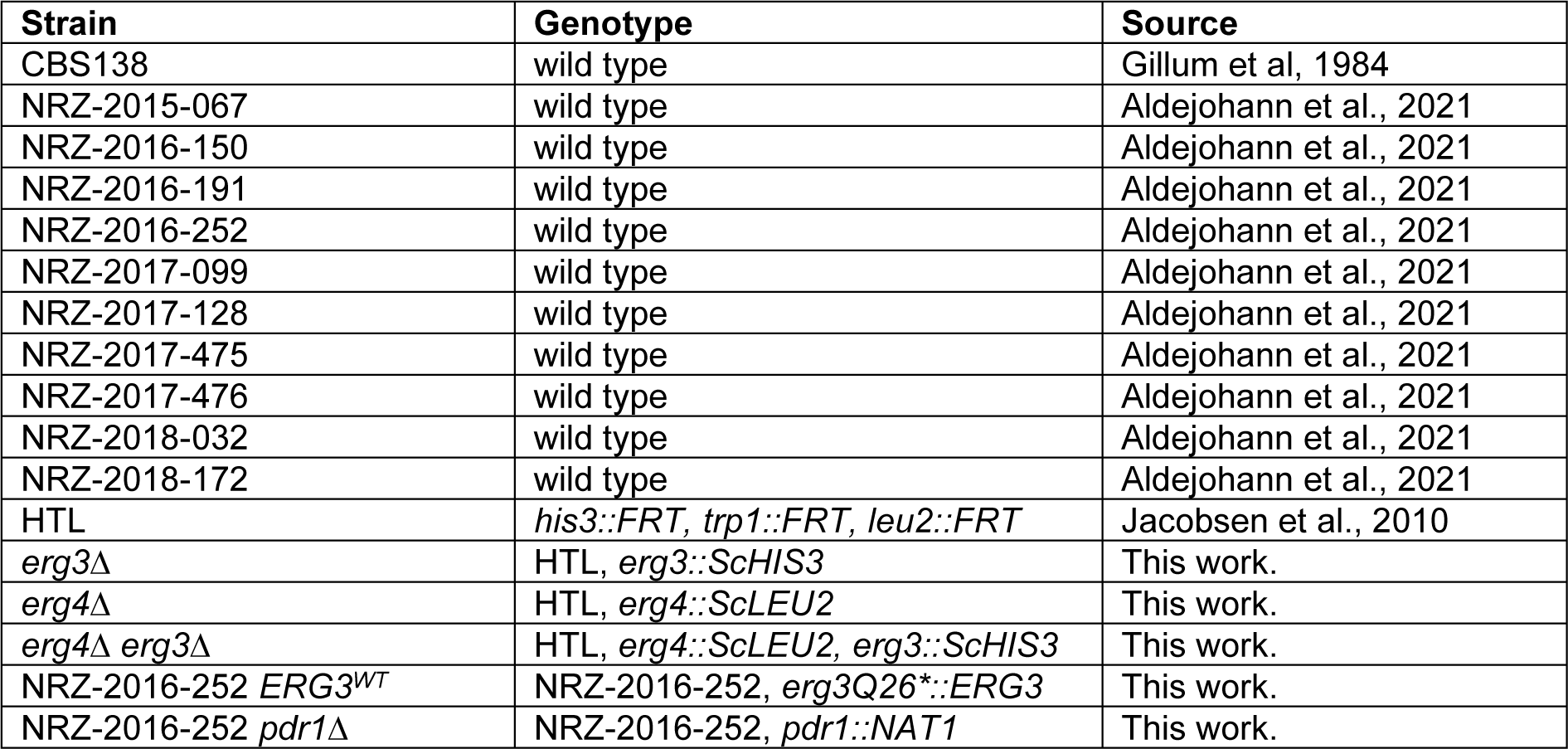
***N. glabratus* strains used in this study.**

**Table S2.**
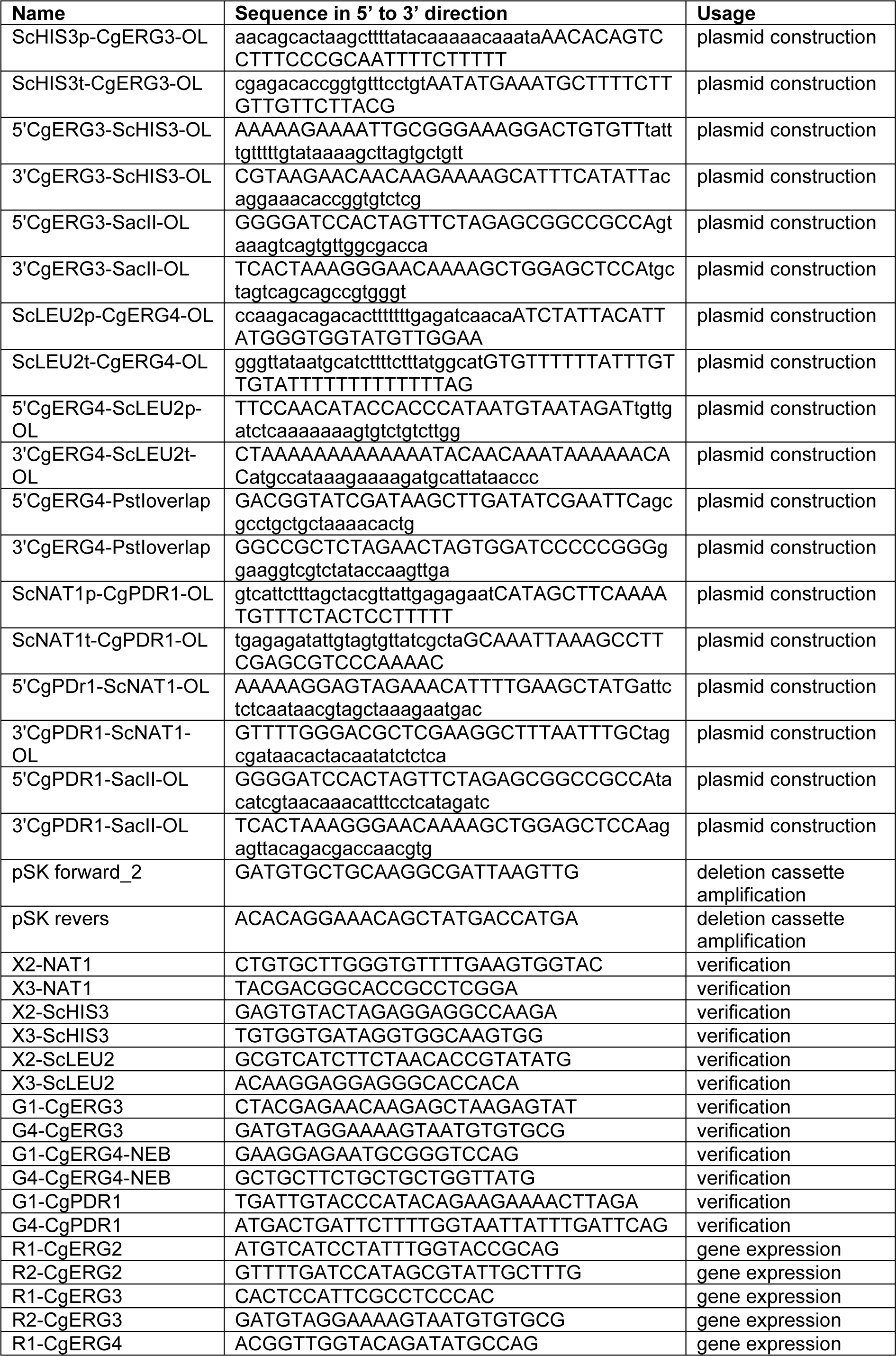
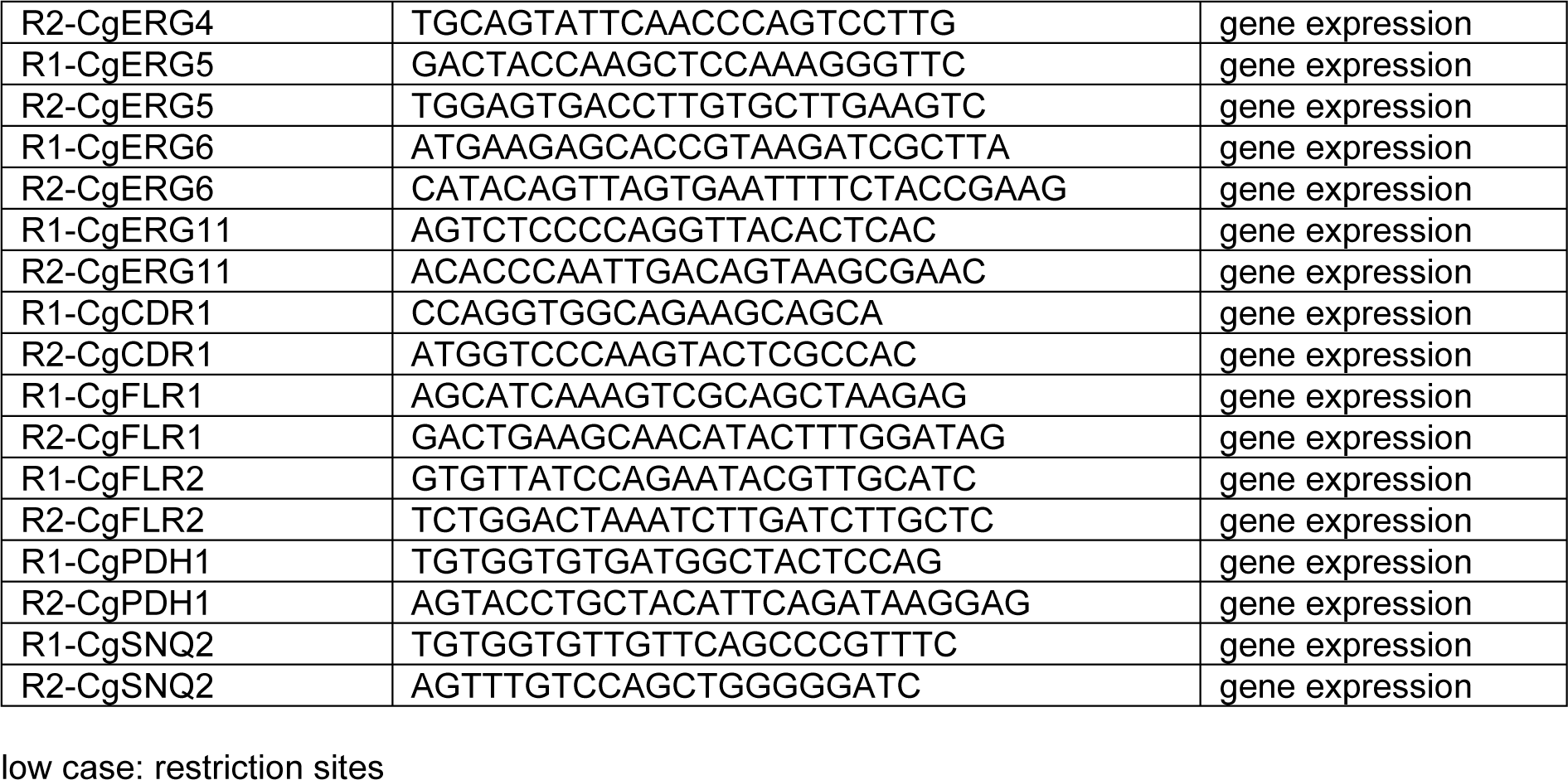
Oligonucleotide primers used in this study.

**Table S3.**
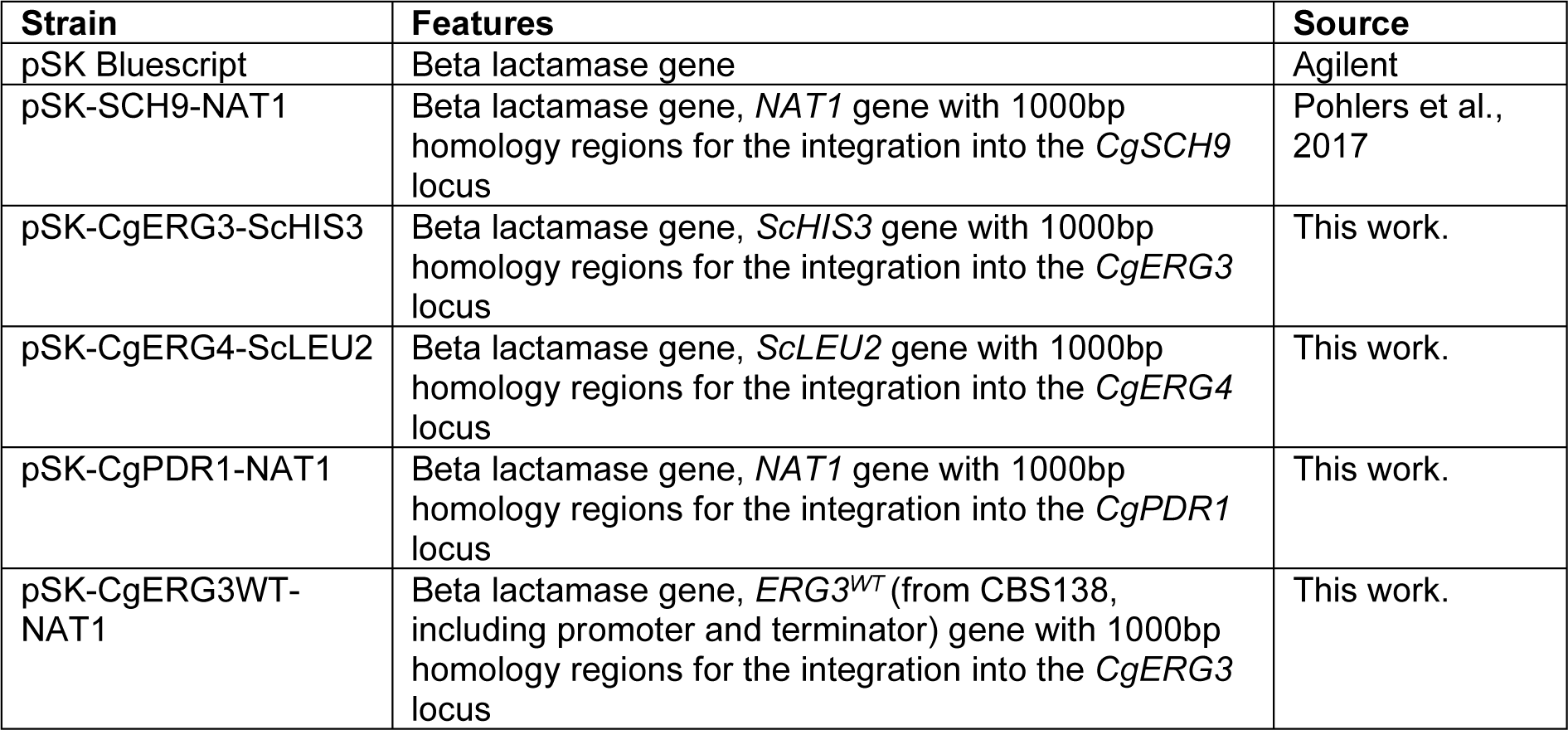
Plasmids used in this study.

