## Supplemental tables S1 to S3 for "Ergosterol-depleted clinical isolates of *Nakaseomyces glabratus* can develop multi-drug resistance without apparent fitness and virulence defects"

**Table S1. *N. glabratus*** **strains used in this study.**

| **Strain** | **Genotype** | **Source** |
| --- | --- | --- |
| CBS138 | wild type | Gillum et al, 1984 |
| NRZ-2015-067 | wild type | Aldejohann et al., 2021 |
| NRZ-2016-150 | wild type | Aldejohann et al., 2021 |
| NRZ-2016-191 | wild type | Aldejohann et al., 2021 |
| NRZ-2016-252 | wild type | Aldejohann et al., 2021 |
| NRZ-2017-099 | wild type | Aldejohann et al., 2021 |
| NRZ-2017-128 | wild type | Aldejohann et al., 2021 |
| NRZ-2017-475 | wild type | Aldejohann et al., 2021 |
| NRZ-2017-476 | wild type | Aldejohann et al., 2021 |
| NRZ-2018-032 | wild type | Aldejohann et al., 2021 |
| NRZ-2018-172 | wild type | Aldejohann et al., 2021 |
| HTL | *his3::FRT, trp1::FRT, leu2::FRT* | Jacobsen et al., 2010 |
| *erg3*∆ | HTL, *erg3::ScHIS3* | This work. |
| *erg4*∆ | HTL, *erg4::ScLEU2* | This work. |
| *erg4*∆ *erg3*∆ | HTL, *erg4::ScLEU2, erg3::ScHIS3* | This work. |
| NRZ-2016-252 *ERG3^WT^* | NRZ-2016-252, *erg3Q26*::ERG3* | This work. |
| NRZ-2016-252 *pdr1*∆ | NRZ-2016-252, *pdr1::NAT1* | This work. |

**Table S2. Oligonucleotide primers used in this study.**

| **Name** | **Sequence in 5’ to 3’ direction** | **Usage** |
| --- | --- | --- |
| ScHIS3p-CgERG3-OL | aacagcactaagcttttatacaaaaacaaataAACACAGTCCTTTCCCGCAATTTTCTTTTT | plasmid construction |
| ScHIS3t-CgERG3-OL | cgagacaccggtgtttcctgtAATATGAAATGCTTTTCTTGTTGTTCTTACG | plasmid construction |
| 5'CgERG3-ScHIS3-OL | AAAAAGAAAATTGCGGGAAAGGACTGTGTTtatttgtttttgtataaaagcttagtgctgtt | plasmid construction |
| 3'CgERG3-ScHIS3-OL | CGTAAGAACAACAAGAAAAGCATTTCATATTacaggaaacaccggtgtctcg | plasmid construction |
| 5'CgERG3-SacII-OL | GGGGATCCACTAGTTCTAGAGCGGCCGCCAgtaaagtcagtgttggcgacca | plasmid construction |
| 3'CgERG3-SacII-OL | TCACTAAAGGGAACAAAAGCTGGAGCTCCAtgctagtcagcagccgtgggt | plasmid construction |
| ScLEU2p-CgERG4-OL | ccaagacagacactttttttgagatcaacaATCTATTACATTATGGGTGGTATGTTGGAA | plasmid construction |
| ScLEU2t-CgERG4-OL | gggttataatgcatcttttctttatggcatGTGTTTTTTATTTGTTGTATTTTTTTTTTTTTAG | plasmid construction |
| 5'CgERG4-ScLEU2p-OL | TTCCAACATACCACCCATAATGTAATAGATtgttgatctcaaaaaaagtgtctgtcttgg | plasmid construction |
| 3'CgERG4-ScLEU2t-OL | CTAAAAAAAAAAAAATACAACAAATAAAAAACACatgccataaagaaaagatgcattataaccc | plasmid construction |
| 5'CgERG4-PstIoverlap | GACGGTATCGATAAGCTTGATATCGAATTCagcgcctgctgctaaaacactg | plasmid construction |
| 3'CgERG4-PstIoverlap | GGCCGCTCTAGAACTAGTGGATCCCCCGGGggaaggtcgtctataccaagttga | plasmid construction |
| ScNAT1p-CgPDR1-OL | gtcattctttagctacgttattgagagaatCATAGCTTCAAAATGTTTCTACTCCTTTTT | plasmid construction |
| ScNAT1t-CgPDR1-OL | tgagagatattgtagtgttatcgctaGCAAATTAAAGCCTTCGAGCGTCCCAAAAC | plasmid construction |
| 5'CgPDr1-ScNAT1-OL | AAAAAGGAGTAGAAACATTTTGAAGCTATGattctctcaataacgtagctaaagaatgac | plasmid construction |
| 3'CgPDR1-ScNAT1-OL | GTTTTGGGACGCTCGAAGGCTTTAATTTGCtagcgataacactacaatatctctca | plasmid construction |
| 5'CgPDR1-SacII-OL | GGGGATCCACTAGTTCTAGAGCGGCCGCCAtacatcgtaacaaacatttcctcatagatc | plasmid construction |
| 3'CgPDR1-SacII-OL | TCACTAAAGGGAACAAAAGCTGGAGCTCCAagagttacagacgaccaacgtg | plasmid construction |
| pSK forward_2 | GATGTGCTGCAAGGCGATTAAGTTG | deletion cassette amplification |
| pSK revers | ACACAGGAAACAGCTATGACCATGA | deletion cassette amplification |
| X2-NAT1 | CTGTGCTTGGGTGTTTTGAAGTGGTAC | verification |
| X3-NAT1 | TACGACGGCACCGCCTCGGA | verification |
| X2-ScHIS3 | GAGTGTACTAGAGGAGGCCAAGA | verification |
| X3-ScHIS3 | TGTGGTGATAGGTGGCAAGTGG | verification |
| X2-ScLEU2 | GCGTCATCTTCTAACACCGTATATG | verification |
| X3-ScLEU2 | ACAAGGAGGAGGGCACCACA | verification |
| G1-CgERG3 | CTACGAGAACAAGAGCTAAGAGTAT | verification |
| G4-CgERG3 | GATGTAGGAAAAGTAATGTGTGCG | verification |
| G1-CgERG4-NEB | GAAGGAGAATGCGGGTCCAG | verification |
| G4-CgERG4-NEB | GCTGCTTCTGCTGCTGGTTATG | verification |
| G1-CgPDR1 | TGATTGTACCCATACAGAAGAAAACTTAGA | verification |
| G4-CgPDR1 | ATGACTGATTCTTTTGGTAATTATTTGATTCAG | verification |
| R1-CgERG2 | ATGTCATCCTATTTGGTACCGCAG | gene expression |
| R2-CgERG2 | GTTTTGATCCATAGCGTATTGCTTTG | gene expression |
| R1-CgERG3 | CACTCCATTCGCCTCCCAC | gene expression |
| R2-CgERG3 | GATGTAGGAAAAGTAATGTGTGCG | gene expression |
| R1-CgERG4 | ACGGTTGGTACAGATATGCCAG | gene expression |
| R2-CgERG4 | TGCAGTATTCAACCCAGTCCTTG | gene expression |
| R1-CgERG5 | GACTACCAAGCTCCAAAGGGTTC | gene expression |
| R2-CgERG5 | TGGAGTGACCTTGTGCTTGAAGTC | gene expression |
| R1-CgERG6 | ATGAAGAGCACCGTAAGATCGCTTA | gene expression |
| R2-CgERG6 | CATACAGTTAGTGAATTTTCTACCGAAG | gene expression |
| R1-CgERG11 | AGTCTCCCCAGGTTACACTCAC | gene expression |
| R2-CgERG11 | ACACCCAATTGACAGTAAGCGAAC | gene expression |
| R1-CgCDR1 | CCAGGTGGCAGAAGCAGCA | gene expression |
| R2-CgCDR1 | ATGGTCCCAAGTACTCGCCAC | gene expression |
| R1-CgFLR1 | AGCATCAAAGTCGCAGCTAAGAG | gene expression |
| R2-CgFLR1 | GACTGAAGCAACATACTTTGGATAG | gene expression |
| R1-CgFLR2 | GTGTTATCCAGAATACGTTGCATC | gene expression |
| R2-CgFLR2 | TCTGGACTAAATCTTGATCTTGCTC | gene expression |
| R1-CgPDH1 | TGTGGTGTGATGGCTACTCCAG | gene expression |
| R2-CgPDH1 | AGTACCTGCTACATTCAGATAAGGAG | gene expression |
| R1-CgSNQ2 | TGTGGTGTTGTTCAGCCCGTTTC | gene expression |
| R2-CgSNQ2 | AGTTTGTCCAGCTGGGGGATC | gene expression |

low case: restriction sites

**Table S3. Plasmids used in this study.**

| **Strain** | **Features** | **Source** |
| --- | --- | --- |
| pSK Bluescript | Beta lactamase gene | Agilent |
| pSK-SCH9-NAT1 | Beta lactamase gene, *NAT1* gene with 1000bp homology regions for the integration into the *CgSCH9* locus | Pohlers et al., 2017 |
| pSK-CgERG3-ScHIS3 | Beta lactamase gene, *ScHIS3* gene with 1000bp homology regions for the integration into the *CgERG3* locus | This work. |
| pSK-CgERG4-ScLEU2 | Beta lactamase gene, *ScLEU2* gene with 1000bp homology regions for the integration into the *CgERG4* locus | This work. |
| pSK-CgPDR1-NAT1 | Beta lactamase gene, *NAT1* gene with 1000bp homology regions for the integration into the *CgPDR1* locus | This work. |
| pSK-CgERG3WT-NAT1 | Beta lactamase gene, *ERG3^WT^* (from CBS138, including promoter and terminator) gene with 1000bp homology regions for the integration into the *CgERG3* locus | This work. |
