## Supplemental material S4. for "Ergosterol-depleted clinical isolates of *Nakaseomyces glabratus* can develop multi-drug resistance without apparent fitness and virulence defects"

**Supplementary Material S4: Extended Methods**

**Stress Tests**

For stress resilience testing, *N. glabratus* strains were grown in YPD medium overnight at 37°C. Out from this preculture, a suspension with 1x 10^8^ fungal cells/mL was made with fresh YPD medium. This suspension was then used to make further dilutions in YPD medium (10^-2^, 10^-4^ and 10^-6^). 5 µl of each dilution was dropped onto YPD agar or YPD plates containing 400 µg/mL Congo Red (Sigma Aldrich, Germany), 1.5 M sodium chloride (Carl Roth, Germany) or 0.0125% SDS (Carl Roth, Germany) and incubated for 3 days at 37°C. Alternatively, YPD plates with drops of fungal cells were incubated at 42°C for 3 days.

**Plasmid construction**

To construct a deletion cassette for *N. glabratus ERG4*, a 1000 bp region upstream of the starting codon was amplified with primers 5’CgERG4-PstIoverlap and 5’CgERG4-ScLEU2p-OL (Table S2). A 1000 bp region downstream of the *CgERG4* stop codon was amplified with primers 3’CgERG4-PstIoverlap and 3’CgERG4-ScLEU2t-OL (Table S2). In addition, the *Saccharomyces cerevisiae* gene *LEU2* was amplified from genomic DNA with primers ScLEU2p-CgERG4-OL and ScLEU2-CgERG4-OL (Table S2). The resulting three PCR products which contained overlapping sequences to each other were then ligated into a *Pst*I-restricted pBluescript (Agilent) by using the NEBuilder HiFi DNA assembly Cloning kit (New England Biolabs, Frankfurt/Main, Germany). The constructed plasmid was named pSK-CgERG4-ScLEU2. The plasmid pSK-CgERG3-ScHIS3 was constructed in a similar way, using the *ScHIS3* gene and a *Sac*II-restricted pSK bluescript. The primers used for plasmid construction are listed in Table S2. For the construction of the *PDR1* deletion cassette, we have amplified the *NAT1* resistance gene from the plasmid pSK-SCH9-NAT1 (Pohlers et al., 2019) and the 1000bp homology regions for integration into the *CgPDR1* locus from genomic CBS138 DNA. Primers used for these amplification steps are listed in table S2. The resulting PCR products were cloned into *Sac*II-restricted pBluescript, leading to plasmid pSK-CgPDR1-NAT1. In a similar way, *ERG3* from CBS138 including promotor and terminator regions was amplified from genomic CBS138 DNA and cloned into *Sac*II-restricted pBluescript, leading to plasmid pSK-CgERG3^WT^-NAT1. The used primers can be found in table S7. All plasmids are listed in Table S3.

**Strain construction**

The deletion cassettes CgPDR1-NAT1, CgERG4-ScLEU2 and CgERG3-ScHIS3 were amplified from the plasmids by using the primers pSK forward-2 and pSK revers (Table S2). For the transformation of the deletion cassettes into *N. glabratus* we have used the lithium acetate protocol as previously described (Gietz et al., 1992). In short, the PDR1-NAT1 cassette was transformed into strain NRZ-2016-252 and transformed cells were plated onto YPD + 100 µg/ml nourseothricin (Jena Bioscience, Jena, Germany) and incubated for two days at 30°C. Transformants were verified with colony PCR using the primer pairs G1-PDR1 / X2-NAT1 and G4-PDR1 / X3-NAT1 to ensure correct integration of *NAT1* into the *CgPDR1* locus (Table S2). The CgERG4-ScLEU2 and CgERG3-ScHIS3 cassettes were transformed into *N. glabratus* HTL which is auxotrophic for histidine, leucine and tryptophane (Jacobsen et al., 2010). Transformed cells were plated onto SDG agar containing required amino acids and incubated for up to three days at 30°C. Transformants were verified with colony PCR and the primers used for verification are listed in Table S2. Consequently, the CgERG3-ScHIS3 cassette was transformed into HTL *erg4::ScLEU2* (*erg4*∆), the transformed cells were plated onto SDG medium with tryptophan and incubated for three days at 30°C. Transformants were verified with colony PCR using primers listed in Table S2.

**Analysis of whole-genome sequencing data**

Paired-end reads were aligned to the *N. glabratus* genome of reference strain CBS138 [GenBank accession GCA_000002545.2] using BWA mem v0.7.12 (Li 2013). Variants were then identified using GATK v4.1.4.1 (Poplin et al., 2017) using the haploid mode and GATK tools MarkIlluminaAdapters, MarkDuplicates, AddOrReplaceReadGroups, HaplotypeCaller for both SNPs and indels, CombineGVCFs, GenotypeGVCFs, GatherVCFs, SelectVariants, and Variant Filtration. Sites were filtered with Variant Filtration using “QD < 2.0 | FS > 60.0 | MQ < 40.0”. Genotypes were filtered if the minimum genotype quality < 50, percent alternate allele <0.8, or depth <10 (Martinez-Zurita and Cuomo, 2023). The variant calling pipeline is detailed in the following github repository: (https://github.com/broadinstitute/fungal-wdl/tree/master/gatk4). Genomic variants were annotated and the functional effect predicted using SnpEff v4.3T (Cingolani et al. 2012). A phylogenetic tree was generated from variant sites that had less than 10% of ambiguous samples using FastTree v2.1.8 (Price et al., 2010).  Counts of unique variants differing between related isolates were obtained using the bcftools view command to select for isolates of interest, followed by bcftools filter with filtering parameters “-i 'AC=1 & AN=2”.

**Cell wall composition analysis**

Staining of the *N. glabratus* cell wall components was performed according to a previously published protocol (Gerwien et al., 2020). We have used an anti-ß-1,3-glucan antibody (1mg/mL, Biosupplies) for the primary staining of ß-1,3-glucan, ConA647 (5mg/mL, Sigma Aldrich) for the primary staining of mannan and WGA-FITC (2mg/mL, Sigma Aldrich) for the primary staining of chitin. Secondary staining was performed with a goat anti-mouse PE-Cy7 antibody (0.2 mg/mL, BioLegend). Flow cytometry was conducted with a CytoFlex (Beckman Coulter) and for each measurement 10,000 events were counted. Data analysis was performed with FlowJo v10.10.0 software. Fluorescence intensities of the clinical isolates were normalized against the fluorescence intensity of control strain CBS138.

**Transcriptome analysis**

RNA quality was checked using a 2100 Bioanalyzer with the RNA 6000 Nano kit (Agilent Technologies). The RIN for all samples was >9.1. cDNA libraries were prepared from 500 ng of total RNA with TruSeq mRNA Stranded Library Prep Kit from Illumina according to manufacturer’s instructions (1/2 volume). Libraries were quantified by QubitTM Flex Fluorometer (ThermoFisher) and quality was checked using 2100 Bioanalyzer with High Sensitivity DNA and DNA 1000 Kit (Agilent). Sequencing of pooled libraries, spiked with 1% PhiX control library, was performed at ~20 million reads/sample in single-end mode with 75 nt read length on the NextSeq 500 platform (Illumina). Demultiplexed FASTQ files were generated with bcl2fastq2 v2.20.0.422 (Illumina).

RNA-seq reads were quality controlled with fastqc v0.11.9 (Andrews, 2010) and adapter clipped with fastp v0.20.0 (Chen et al., 2018). After another quality control, reads were mapped to the reference genome ASM254v2 using the STAR v2.7.10 tool (Dobin et al., 2013). Annotation count files were generated with featureCounts v2.0.6 (Liao et al., 2014). Gene-based analysis was performed with DESeq2 v1.38.3 filtering out variants with p_adj_ ≤ 0.05 and an absolute fold change ≥ 2 for upregulated genes and an absolute fold change ≤0,5 for downregulated genes (Love et al., 2014). The Volcano Plot was prepared with the R package EnhancedVolcano v1.16.0. Heatmap plots were prepared with the R package pheatmap v1.0.12.
